# A simple Ca^2+^-imaging approach to neural network analysis in cultured neurons

**DOI:** 10.1101/2020.08.09.243576

**Authors:** Zijun Sun, Thomas C. Südhof

## Abstract

**Background:** Ca^2+^-imaging is a powerful tool to measure neuronal dynamics and network activity. To monitor network-level changes in cultured neurons, neuronal activity is often evoked by electrical or optogenetic stimulation and assessed using multi-electrode arrays or sophisticated imaging. Although such approaches allow detailed network analyses, multi-electrode arrays lack single-cell precision, whereas optical physiology generally requires advanced instrumentation.

**New Method:** Here we developed a simple, stimulation-free protocol with associated Matlab algorithms that enables scalable analyses of network activity in cultured human and mouse neurons. The approach allows analysis of overall networks and single-neuron dynamics, and is amenable to scale-up for screening purposes.

**Results:** We validated the protocol by assessing human neurons with a heterozygous conditional deletion of Munc18-1, and mouse neurons with a homozygous conditional deletion of neurexins. The approach described here enabled identification of differential changes in these mutant neurons at the network level and of the amplitude and frequency of calcium peaks at the single-neuron level. These results demonstrate the utility of the approach.

**Comparison with existing method:** Compared with current imaging platforms, our method is simple, scalable, and easy to implement. It enables quantification of more detailed parameters than multi-electrode arrays, but does not have the resolution and depth of more sophisticated yet labour-intensive analysis methods, such as electrophysiology.

**Conclusion:** This method is scalable for a rapid assessment of neuronal function in culture, and can be applied to both human and mouse neurons. Thus, the method can serve as a basis for phenotypical analysis of mutations and for drug discovery efforts.

## 1. INTRODUCTION

Ca^2+^-imaging provides a wide range of applications in neuroscience because it allows monitoring large populations of neurons *in vivo* and in culture^1^. In culture, neurons exhibit two primary activity modes, sparse or burst spiking of individual neurons and synchronous repetitive firing of multiple neurons in networks. Depending on the developmental stage of cultured neurons, Ca^2+^-imaging typically reveals some spontaneous firing of neurons^2–4^. As neurons form mature neural networks, synchronous firing of neurons via network activity increases in parallel with the synapse density, and can be visualized by Ca^2+^-imaging^4^. The properties of network activity in cultured neurons depends not only on the extent of synaptic communication between neurons, but also on the electrical properties and the axonal and dendritic development of these neurons as well as on their cellular state^2–5^. Therefore, network firing of neurons represents a proxy for a broad range of neuronal properties, including their developmental maturity, synaptic connectivity, and signalling status.

A challenge in measuring the network activity of cultured neurons is that synchronicity occurs stochastically and depends on culture quality (density, age etc)^3,6^. As a result, induction of network activity often requires external stimulation, such as using electrical stimulation or optogenetics to induce responses^7^. Recently, imaging-based platforms have been developed for sophisticated all-optical ‘electrophysiology’, which allow network-level analyses by applying optical stimuli and monitoring individual cells via optical reporters^8,9^. A major advantage of external stimulation in network analyses is that it enables observation of how synchronized firing is induced, i.e., provides causality. A disadvantage is that the delivery of stimuli introduces additional complexity to the imaging setup, for example implementation of electrodes or installation of optical lightguides for stimulation. Moreover, variations in the stimulus efficacy may produce variable responses^10^. An alternative approach to stimulating network activity in cultured neurons is the application of an activating pharmacological agent. This approach is particularly useful for neurons derived from human embryonic stem (ES) or induced pluripotent stem (iPS) cells that may be difficult to analyse because they are generally less mature than neurons in primary cultures and exhibit less spontaneous network activity^8^. However, pharmacologically induced responses may differ from more physiological spontaneous responses^11^.

The abovementioned issues underscore the need for less sophisticated Ca^2+^-imaging approaches that enable rapid analyses of sparse and synchronous neuronal activity and facilitate studies of stem cell-derived neurons. Here, we describe such an approach. We describe a simple Ca^2+^-imaging protocol that induces spontaneous network activity in human and mouse neurons without using external stimulation. Moreover, we provide an analysis pipeline that quantifies network activity and single-neuron dynamics. We offer procedures for the culture and imaging of human and mouse neurons, step-wise illustrations of the analysis pipeline, and protocols for the interpretations of the results. Network activity of cultured human neurons was robustly induced in the optimised Ca^2+^-imaging conditions, rendering it suitable for the study of disease-associated mutations in neurons. We tested the sensitivity and robustness of this approach using mutants that were previously shown to exhibit defects in synaptic transmission^12,13^, and validated our results using established calcium analysis software programs.

Ca^2+^-imaging monitors intracellular Ca^2+^-fluxes that are induced by neuronal depolarization and action potential firing^14^. Ca^2+^-fluxes are generally monitored via the fluorescence signal of GCaMP-type Ca^2+^-indicators, which needs to be analysed to extract the features of individual Ca^2+^ spikes and to infer the underlying neuronal activity^1^. The image analysis algorithms we wrote quantifies multiple parameters, including the synchronicity rate of the network activity and the Ca^2+^-spike dynamics of individual neurons, including their amplitude and frequency. The goal of the present protocol is to establish an easily accessible and reproducible Ca^2+^-imaging approach that can be scaled up for mutagenesis or drug screening purposes.

## 2. Materials and Methods

### 2.1 Generation of Ngn2-induced human neuron expressing GCaMP Ca^2+^-indicators

We used a co-culture system of induced human neurons (iN cells) with mouse glia, which provides a supporting layer and factors required for neuronal survival, dendritic arborization, and synapse formation^15–17^ (**Fig. 1A**). Human neurons were trans-differentiated from H1 ES cells that were cultured on Matrigel-coated 6-well plate according to the manufacturers’ protocols (Corning), and maintained in culture medium containing mTeSR plus basal medium and 5 x supplement (Stem Cell Technologies). Thiazovivin (2 μM, BioVision) was supplemented into the seeding media. Induction of iN cells was performed via lentiviral delivery of the tetracycline transactivator (rtTA) and tetracycline-responsive (*TetO*)-driving Ngn2 as described^15^. H1 ES cells were infected at the same time with lentiviruses expressing GCaMP6m under control of the human synapsin-1 promoter. Lentiviruses were produced in HEK293T cells by calcium phosphate transfection^18^. After lentiviral transduction, cells were maintained in DMEM/F12 medium supplemented with BDNF (10 ng/ml, PeproTech), laminin (0.2 μg/ml, Invitrogen), human NT-3 (10 ng/ml, PeproTech), N2 supplement (ThermoFisher Scientific), Doxycycline (2 μg/ml, Sigma-Aldrich), and non-essential amino acid solution (ThermoFisher Scientific) with daily media changes. Puromycin (1 μg/ml, InvivoGen) was added for two days from DIV2. On DIV4, the developing human iN cells were dissociated with Accutase (Innovative Cell Technologies) and seeded at a density of 1.8-2×10^5^ cells/coverslip onto overnight Matrigel-coated coverslips (in 24-well plates) containing mouse glia that had been plated 24-48 hrs previously. After DIV4, the mixed neuron/glia cultures were maintained in Neurobasal medium (ThermoFisher Scientific) supplemented with B27 Supplement (ThermoFisher Scientific) and Glutamax (ThermoFisher Scientific), with a weekly change of half of the medium. On DIV6, Ara-C (Sigma-Aldrich) was added to the culture medium (2 μM final concentration). On DIV10, fetal bovine serum (ATLANTA Biological; final concentration = 2-3%) were added to the culture medium. Ca^2+^-imaging was performed at DIV35-40 (**Fig. 1A**).

**Fig. 1.**
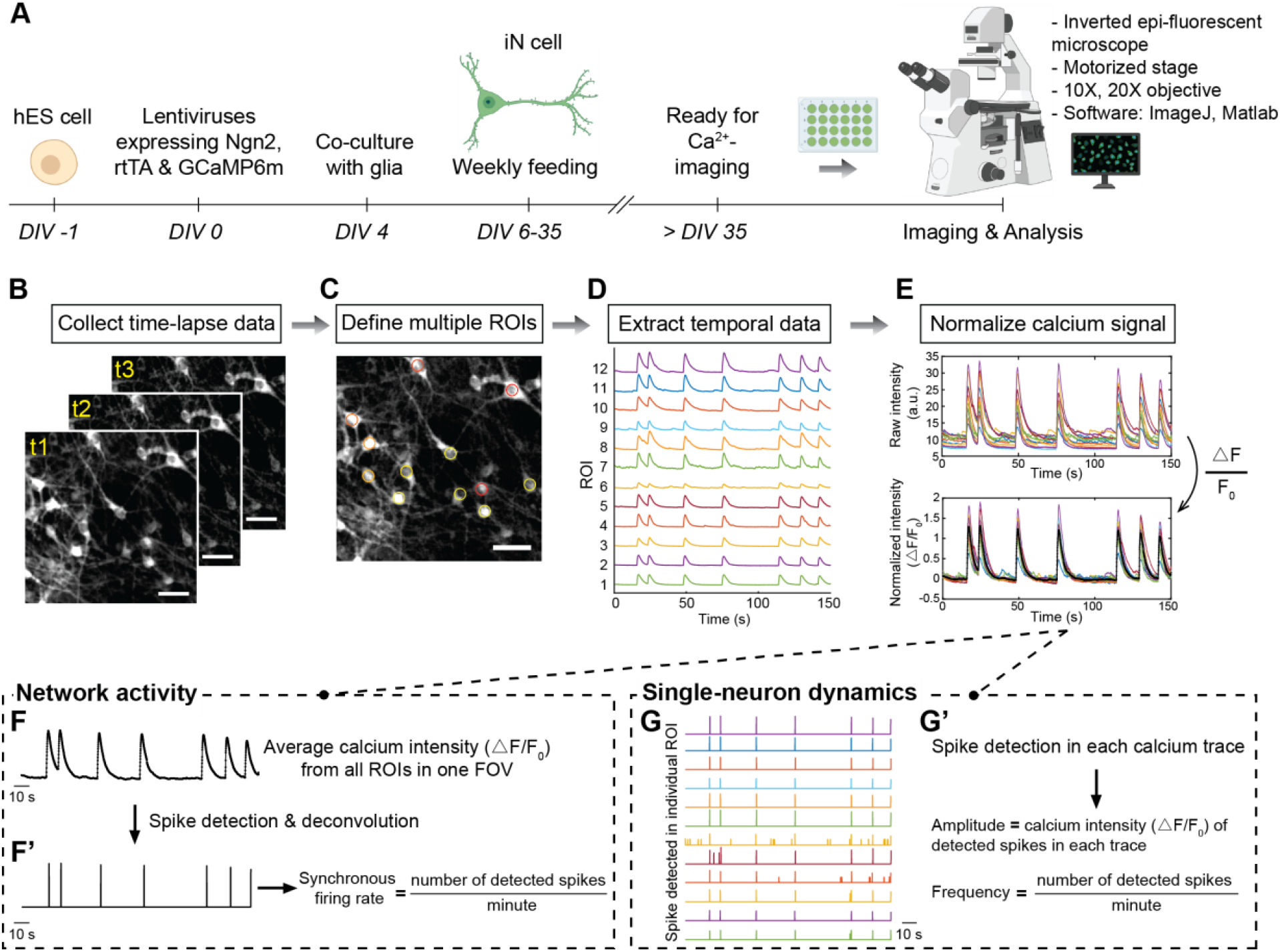
Ca^2+^-imaging and analysis workflow. (**A**) Experimental strategy for culturing and imaging induced human neurons (iN cells). At 0 days *in vitro* (DIV0), human ES or iPS cells were infected with lentiviruses expressing Ngn2 (from the tetO promoter activated by co-expressed rtTA) and with lentiviruses expressing GCaMP6m (under the synapsin promoter). At DIV4, cells were co-cultured with mouse glia. Cells were maintained by weekly feeding up to DIV35 for maturation of Ca^2+^-imaging. For image acquisition and analysis, an inverted epi-fluorescent microscope with a digital or CCD camera and a 10x or 20x objective connected to a PC or Macintosh equipped with the image analysis software ImageJ and with Matlab are required. A motorized microscope stage with robotic timing of imaging is optional. (**B-E**) Basic image processing pipeline. (**B**) Example of time-lapse images (yellow texts “t1, t1, t3” indicate time series). Images are ‘stacked’ to provide a maximal intensity projection (**C**). (**C**) Maximal intensity projected image superimposed with ROI boxes (coloured circles) at neuron soma. (**D**) Raw intensity of the Ca^2+^ signal are extracted from selected ROIs. (**E**) Normalization of calcium signal. Coloured lines in both panels indicate individual Ca^2+^ traces and the black thick line in the lower panel indicates the average intensity from all traces. (**F & F’**) Illustration of network activity analysis. (**F**) The average intensity is calculated from Ca^2+^ traces in all ROIs in one FOV (black thick line in lower panel in **E**) and is subsequently used for spike detection and deconvolution using a built-in Matlab spike-detection function. (**F’**) Illustration of quantification of synchronous firing rate. Synchronous spikes are identified using a threshold based on predefined percentile (mean + 1.5-2 X standard deviation, s.d.). Synchronicity rate is quantified as the number of the detected synchronous spikes in one minute. (**G-G’**) Illustration figure of the quantification of single-neuron dynamics. (**G**) Example of spikes detected in individual Ca^2+^ traces. The spike detection and deconvolution is performed using the same method as illustrated in panel **F’**. (**G’**) Quantification of single-neuron amplitude and frequency. The amplitude of peaks in each trace is defined as the mean value of ΔF/F_0_ of individual peak. The frequency of peaks in each trace is defined as the number of detected peaks in one minute. Scale bars, 25 μm. Objective, 20 X.

### 2.2 Generation of mouse cortical primary neuron with GCaMP expression

Cortical primary neurons from newborn triple neurexin-1/2/3 conditional KO mice^13^ were seeded on Matrigel-coated coverslips in a 24-well plate in MEM medium (ThermoFisher Scientific). On DIV2, the medium was changed to Neurobasal medium containing 2 μM AraC. On DIV3, half of the medium was exchanged, and the cells were infected with two lentiviruses that encoded (1) GCaMP6m and (2) ΔCre-P2A-mCherry or Cre-P2A-mCherry (all driven by the human synapsin promoter). Cultures were maintained through DIV14-16, when Ca^2+^-imaging was performed.

### 2.3 Ca^2+^-imaging

Coverslips were gently washed twice in modified Tyrode solution (25 mM HEPES (Invitrogen), 140 mM NaCl, 5 mM KCl, 1 mM MgCl_2_, 10 mM glucose, 2 mM CaCl_2_, 10 μM glycine pH 7.2–7.4, pre-warmed to 37 °C), and placed into a glass-bottom 12-well plate (Cellvis) containing Ca^2+^-imaging buffer (25 mM HEPES, 140 mM NaCl, 8 mM KCl, 1 mM MgCl_2_, 10 mM glucose, 4 mM CaCl_2_, 10 μM glycine pH 7.2–7.4, pre-warmed to 37 °C). After 1-2 min equilibration, Ca^2+^-imaging was performed on an inverted epi-fluorescence microscope (Nikon EclipseTS2R, DS-Qi2 digital camera) with the 488 nm filter at room temperature. GCaMP6m fluorescence was recorded for 2-3 min at a frame rate of 4-10 frames/s. Approximately 800-3000 time-lapse images depending on the frame rate (536×536 pixel resolution, 14-bit grayscale depth for human and mouse cultures) were acquired at a 1 x digital zoom using either a 10x or 20x objective. Acquisition time of each frame can vary between 0.01-0.1s, depending on the camera used for detection and the exposure time subjected to the brightness of GCaMP expression; long-term recording can cause photobleaching and phototoxicity. Per coverslip, 2-3 fields were imaged and a minimal of 3 coverslips were recorded for each biological batch. For each batch, all images were acquired using the same light intensity and exposure time. Time-lapse imaging was performed in a field of view (FOV) containing confluent neuron populations with non-overlapping soma.

### 2.4 Image pre-processing and network and spike analysis algorithms

Time-lapse image files were converted to *.tiff /.tif* format using ImageJ (https://imagej.nih.gov/ij/). Quantifications were performed using home-written Matlab functions and scripts included in **Appendix A** and **B**. Datasheets containing raw Ca^2+^ traces are used as inputs for all algorithms. The datasheets should be organised such that rows correspond to times and columns to individual cells. To determine the network activity of the neuronal culture based on the synchronous firing rate of the entire cell population in the FOV, follow section 3.2 (**Appendix B3**); to determine the single-neuron activity in terms of amplitude and frequency, follow section 3.3 (**Appendix B4**). An example for the variable definition and the output can be found in the instruction (**Appendix A**). For measuring network activity, the algorithm in **Appendix B3** provides the quantification of synchronous firing rate and synchronous peak amplitude as two outputs, and we only used the former for illustration here.

### 2.5 Statistical analyses

Matlab (MathWorks) was used to process images and quantifications from time-lapse tiff files. Data were exported from Matlab to .*csv* files which were imported for plotting into R software (http://www.r-project.org, version 1.2.5001; R packages ggplot2, tidyverse, lubridate). The boxplots were plotted using the R package geom_boxplot, ggplot2 0.9.3.1. The upper and lower hinges of the box indicate the 25th and 75th percentiles. The upper whisker and the lower whisker extend from the hinge of the box to the highest and lowest value that is within 1.5 X interquartile of the hinge, respectively. All *P* values were calculated using Student’s *t*-test with *P* < 0.05 considered significant, unless otherwise stated.

## 3. Results

### 3.1 Imaging of neurons and image analysis

Human neurons expressing GCaMP6m and co-cultured with mouse glia were imaged in a standard epi-fluorescence microscope, and the GCaMP6m fluorescence signal was recorded as a function of time (**Fig. 1A**). Regions of interests (ROIs) containing the soma region of neurons to be analysed were manually selected using the Matlab algorithm provided in **Appendix B1**. The algorithm first projected the time-lapse stack to its maximum intensity image to allow a user to find and select a neuron (**Fig. 1B**). Upon selection of the center of the soma of a neuron, a circular ROI mask was superimposed onto the selected neuron (**Fig. 1C**). The size of the ROI can be specified in the algorithm according to the acquisition resolution (see **Appendix A** for detailed instructions and examples of variable definitions). Each ROI was defined as a node to extract the raw Ca^2+^ intensity from the ROI, which was used for all further analysis. After selection of all desired ROIs, traces of the GCaMP6m fluorescence intensity per ROI over time were generated (**Fig. 1D**). The intensity profile was automatically saved into an *.mat* file named after the image name in a folder automatically created. The raw data is in the matrix *ROI_intensity1*, in which each row corresponds to a single frame and each column corresponds to an ROI and can be found by loading the *.mat* file in the command window. To plot the raw intensity of the Ca^2+^ signal, load the *ROI_intensity1* matrix generated from the above step and run the script in **Appendix B2**. This will return a figure showing all traces (upper panel in **Fig. 1E**). The baseline fluorescence level (F_0_) of each trace was calculated as the average basal fluorescence intensity during the first, middle and last 100 frames of each trace^12,19^. The Ca^2+^ signal intensity of each raw trace was then normalized by calculating ΔF/F_0_ as the ratio of the increase in fluorescence (ΔF) to the baseline (F_0_) (lower panel in **Fig. 1E**).

### 3.2 Analysis of network activity by inducing synchronous activity

In cultured Ngn2-induced neurons trans-differentiated from ES cells, individual neurons exhibited occasional spontaneous Ca^2+^-spikes (**Fig. 2A-A’**), but synchronous firing of two or more neurons was rare (**Fig. 2A-A’, C**). Therefore we elevated neuronal firing by increasing the concentrations of Ca^2+^ and K^+^ in the imaging solution to 4 mM and 8 mM, respectively. Under this condition, frequent synchronous bursting events were observed in cultured neurons (**Fig. 2B-B’, D**). To quantify the synchronous firing activity, we analysed network-wide synchronous Ca^2+^ spikes. First, we obtained the global intensity profile of Ca^2+^ transients by calculating the average intensity of Ca^2+^ signal from all ROIs in one FOV (thick black line in lower panel in **Fig. 1E, Fig. 1F**). Second, we used a built-in Matlab function for spike detection and identified the spikes based on the average profile above the threshold (here we used mean + 1.5-2 x s.d.). The synchronous firing rate was defined as the number of the synchronous Ca^2+^ spikes per minute (**Fig. 1F’**). The synchronicity rate and the amplitude of synchronous spikes was quantified by the script in **Appendix B3**. Here, we only include the synchronicity rate for discussion and illustration purposes. We then compared the network activity of the culture in normal Tyrode medium versus the imaging solution that contains elevated Ca^2+^ (4 mM) and K^+^ (8 mM) concentrations. We found that the synchronous firing rate was significantly higher in the imaging solution than in Tyrode medium (**Fig. 2G**). In order to understand the variability of the Ca^2+^ signals observed in cultured human neurons (iN cells), we quantified the coefficient of variation of the synchronous firing rate (defined as the ratio of the standard deviation of the synchronous firing rate to its mean value (i.e., s.d. / mean in a given FOV). Comparison of the coefficient of variation between neurons cultured in normal Tyrode medium versus the imaging solution did not uncover a significant difference (**Fig. 2G’**), suggesting that the network activity measured by synchronous firing rate is comparable across iN cell cultures. Ngn2-induced human neurons are composed almost exclusively of excitatory neurons^15^. As a result, addition of CNQX (10 μM), an antagonist of AMPA-type glutamate receptors, suppressed the network activity of human neurons, although isolated Ca^2+^ spikes could still be observed (**Fig. 2E**). Treatment of the cultured neurons with tetrodotoxin (TTX; 1 μM), a Na^+^-channel blocker, also abolished synchronous network activity of neurons, as evidenced by a complete loss of Ca^2+^ spikes, which would be expected for network activity driven by action potential firing (**Fig. 2F**). Note that TTX also ablates sparse spontaneous Ca^2+^ spikes, supporting the notion that spontaneous non-synchronous Ca^2+^ spikes also depend on action potential firing (**Fig. 2F**).

**Fig. 2.**
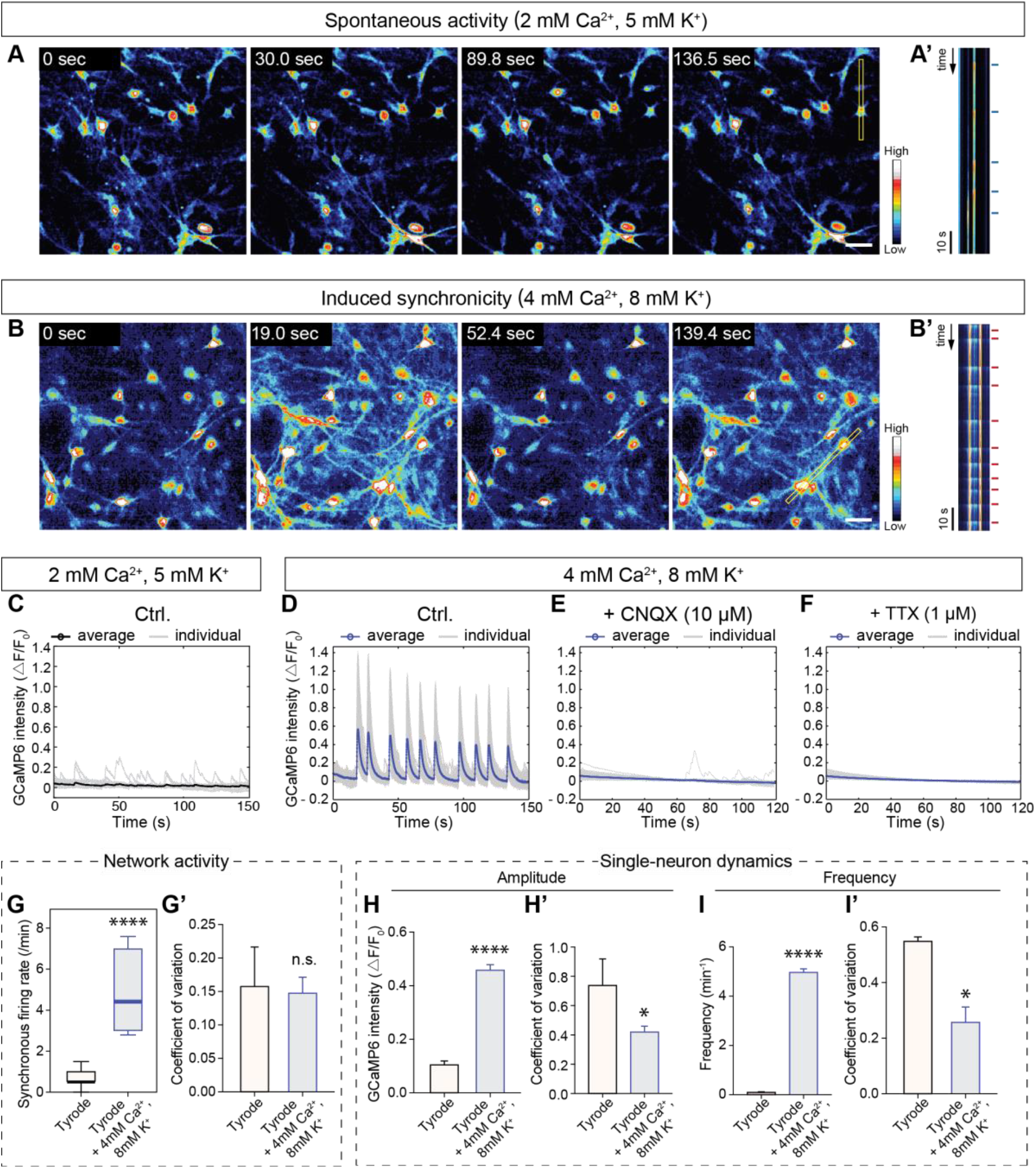
Illustration of the analysis of network activity and single-neuron dynamics of cultured human neurons. (**A** & **A’**) Representative time-lapse images of the spontaneous activity of human neurons expressing GCaMP6m monitored in Tyrode solution (**A**). The kymograph (**A’**; represents two neurons are from the yellow box in the rightmost panel in **A**) for the first 90 s of recording showed non-synchronous spikes, as indicated by blue lines. (**B & B’**) Same as **A & A’**, but for neurons examined in modified Tyrode solution with an increased Ca^2+^ (4 mM) and K^+^ concentration (8 mM). In the kymograph (**B’**), synchronous spikes are indicated by red lines. (**C**) Ca^2+^ signal traces plotted from (**A**). No significant synchronous peaks are observed from the averaged intensity (black line), but individual Ca^2+^ spikes are present (light-grey lines). (**D**) Ca^2+^ traces plotted from (**B**). Synchronous peaks are observed in the averaged intensity (blue line; light-grey lines indicate individual Ca^2+^ traces). (**E & F**) Representative Ca^2+^ traces monitored in neurons in modified Tyrode solution with an increased Ca^2+^ and K^+^ concentration and in the presence of 10 μM CNQX (**E**) or 1 μM TTX (**F**). (**G & G’**) Quantification of network activity of cultures in normal Tyrode and that in 4 mM Ca^2+^, 8 mM K^+^ condition using the synchronicity rate from this study (**G**) and the coefficient of variation of synchronous firing rate (**G’**). (**H & H’**) Quantification of the single-neuron amplitude (**H**) and coefficient of variation of the amplitude (**H’**) of cultures in normal and modified Tyrode solutions. (**I & I’**) Quantification of the single-neuron frequency (**I**) and coefficient of variation of the frequency (**I’**) of cultures in normal and modified Tyrode solutions. Data in **G-I’** represent mean ± SEM. In **G-I’**, n = 146 and 216 neurons were measured for the normal and modified Tyrode solution conditions. Student’s *t*-tests were used for all groups (*, p<0.05; ****, p<0.0001; n.s., not significant). Scale bars, 25 μm. Objective, 20 X.

### 3.3 Quantification of single-neuron amplitude and frequency

To analyse the single-neuron Ca^2+^ dynamics, we computed the amplitude and frequency, which are common parameters quantified in Ca^2+^ imaging analyses (**Fig. 1G-G’**). We used the spike detection method described above, using a threshold of the mean + 1.5-2.0 x s.d. in each Ca^2+^ trace^12,19^ (**Fig. 1G**). The single-neuron amplitude was calculated from all detected spikes within an individual Ca^2+^ trace after normalization (*ΔF/F_0_*, **Fig. 1G’**)^1^. The single-neuron frequency was calculated as the number of detected spikes in each Ca^2+^ trace per minute (**Fig. 1G’**). The quantification program is provided by script in **Appendix B4**. Running the program produces two graphs: a plot of the detected spikes (example in **Fig. 1G**) and a plot of the Ca^2+^ traces after normalization (example in lower panel in **Fig. 1E**). We compared the single-neuron amplitude and frequency of cultures in normal and modified Tyrode solution. We found that the amplitude and frequency were both increased by the 4 mM Ca^2+^ and 8 mM K^+^ concentrations in the modified Tyrode solution as expected (**Fig. 2H & I**). The neuron-to-neuron variation was quantified as the coefficient of variation of the amplitude and frequency (i.e., s.d. / mean per neuron), revealing that the coefficient of variation of both decreased in the modified Tyrode solution as expected (**Fig. 2H’ & I’**).

### 3.4 Validation of the proposed Ca^2+^ imaging protocol by analysis of human neurons with a conditional Munc18-1 (STXBP1) deletion

To validate our Ca^2+^ imaging approach, we analysed a previously characterized mutation affecting synaptic transmission, the heterozygous conditional knockout (cKO) of Munc18-1 (human gene symbol: *STXBP1*)^20^. Munc18-1 is a central component of the synaptic vesicle fusion machinery (reviewed in reference^21^), and heterozygous Munc18-1 deletions cause a severe neurodevelopmental syndrome in human infants called Ohtahara syndrome. We converted human H1 ES cells carrying the conditional Munc18-1 deletion into neurons using Ngn2 expression, and expressed in these neuron either an inactive mutant Cre (ΔCre, control) or active Cre (to induce the Munc18-1 deletion) in addition to GCaMP6m. Munc18-1 cKO H1 cells were co-infected on DIV0 with lentiviruses expressing Cre-P2A-mCherry or ΔCre-P2A-mCherry and with lentiviruses expressing Ngn2, rtTA and GCaMP6m (**Fig. 3B & C**). Immunoblotting confirmed that Cre-expressing Munc18-1 cKO neurons exhibited a large reduction in Munc18-1 protein levels (**Fig. 3A**). We performed Ca^2+^ imaging on the resulting neurons at DIV35-40, which revealed a dramatic change in Ca^2+^ spiking and synchronous network activity in Munc18-1 mutant neurons (**Fig. 3B’ & C’**).

**Fig. 3.**
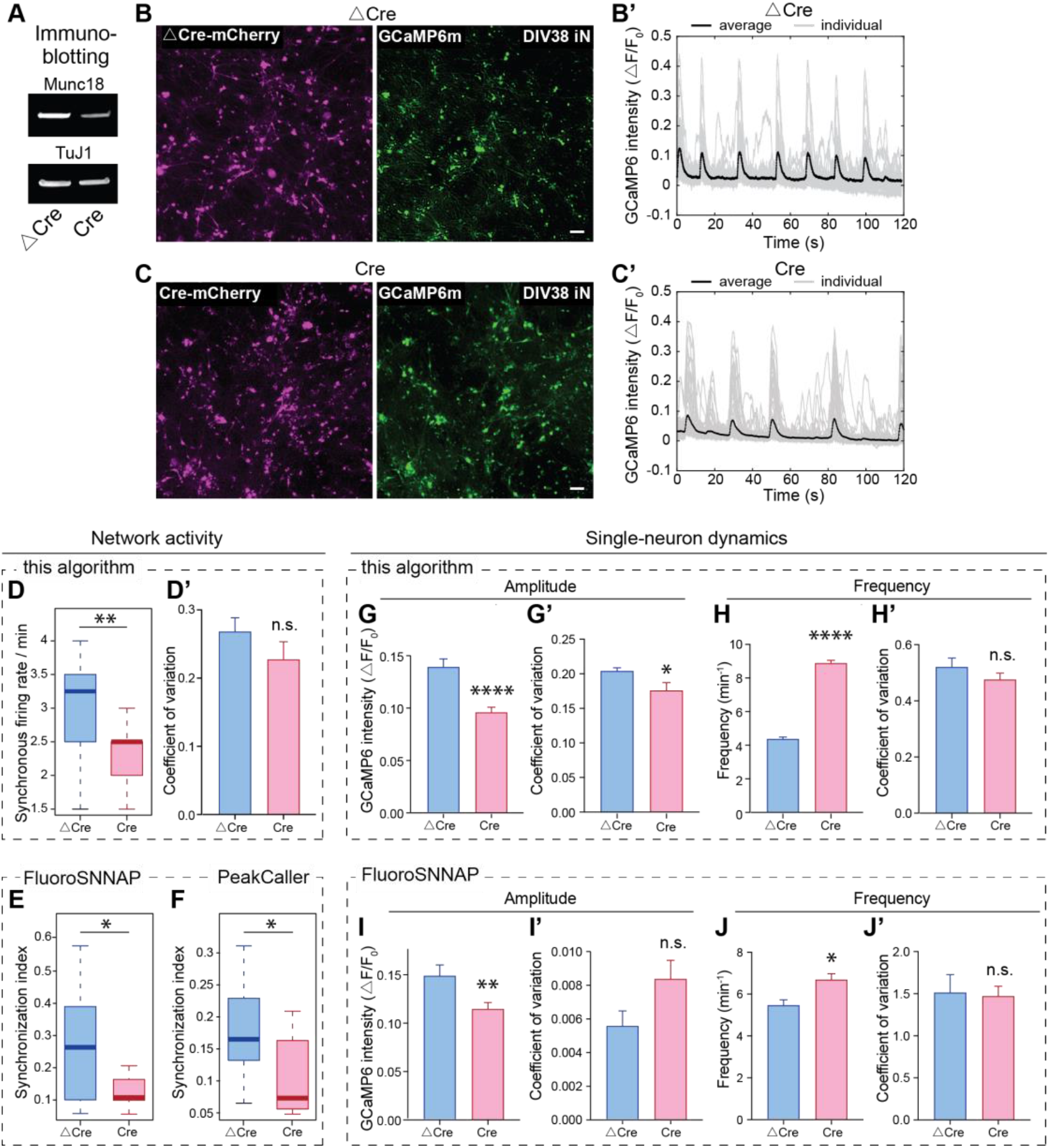
Analysis of network and single-neuron activities in Munc18-1 cKO human neurons. (**A**) Immunoblots of control (ΔCre) and heterozygous mutant neurons (Cre). (**B & C**) Representative images of the mCherry (left) and the GCaMP6m signal (right) observed in heterozygous human Munc18-1 cKO neurons expressing ΔCre (control) or Cre (deletion) as analysed at DIV38. (**B’** & **C’**) Individual (thin grey lines) and averaged Ca2+ signal traces (thick black lines) of heterozygous human Munc18-1 cKO neurons expressing ΔCre (control, **B’**) or Cre (deletion, **C’**) as recorded at DIV38. (**D & D’**) Quantification of network activity using synchronicity firing rate (**D**) and the coefficient of variation of synchronous firing rate (**D’**) for control and heterozygous Munc18-1mutant neurons. (**E & F**) Quantification of synchronization index by FluoroSNNAP (**E**) and PeakCaller (**F**) for control and heterozygous mutant neurons. In FluoroSNNAP, the synchronization index was quantified by separating the time-varying fluorescence signals into its amplitude and phase components and comparing the similarity of timing of events (phase) of all the ROIs. In PeakCaller, the synchronization index was computed pairwise based on autocorrelation function produced from each trace and the synchronization index was calculated from the autocorrelation functions from all input ROIs. (**G & G’**) Quantification of single-neuron amplitude (**G**) and the coefficient of variation of amplitude (**G’**) for control and heterozygous Munc18-1 mutant neurons using program in this study. (**H & H’**) Quantification of single-neuron frequency (**H**) and the coefficient of variation of frequency (**G’**) for control and heterozygous Munc18-1 mutant neurons using program in this study. (**I & I’**) Quantification of single-neuron amplitude (**I**) and the coefficient of variation of amplitude (**I’**) for control and heterozygous Munc18-1 mutant neurons using FluoroSNNAP. (**J & J’**) Quantification of single-neuron frequency (**H**) and the coefficient of variation of frequency (**G’**) for control and heterozygous Munc18-1 mutant neurons using FluoroSNNAP. 689 and 981 neurons from control and Munc18-1-mutant neurons from 3 biological replicates were used in **D-D’, F-H’**. 1139 and 1274 detected ROIs from control and Munc18-1-mutant neurons from 3 biological replicates were used in **E, I-J’**. 8 FOVs from 3 biological batches were recorded for control and mutant human neurons each. Data in **D-J’** represent mean ± SEM. Student’s *t*-tests were used for all groups (*, p<0.05; **, p<0.01; ****, p<0.0001; n.s., not significant). Scale bars, 100 μm. Objective, 10 X.

Analysis of the neuronal Ca^2+^ spikes demonstrated that the heterozygous Munc18-1 deletion caused a ~50% suppression of neuronal network activity, which was quantified as the synchronous firing rate (**Fig. 3D**). At the single-neuron level, the Munc18-1 deletion produced a ~30% decrease in Ca^2+^ signal amplitude (**Fig. 3G**) and a significant increase in Ca^2+^ signal frequency (**Fig. 3H**). It was noteworthy that the increased frequency of Ca^2+^ spikes exhibited in Munc18-1 mutant neurons are asynchronous (**Fig. 3C’**). The increase in Ca^2+^ signal frequency despite a decrease in network activity is due to the emergence of aberrant Ca^2+^ spikes in the mutant neurons upon suppression of synchronous firing, consistent with the impairment in synaptic communication induced by the Munc18-1 deletion. Moreover, we observed a modest decrease of the coefficient of variation of the single-neuron amplitude in the Munc18-1 mutant neurons compared with the control, presumably because the Ca^2+^ spikes are reliably smaller in amplitude (**Fig. 3G’**), No significant difference in the coefficient of variation of the synchronous firing rate (**Fig. 3D’**) and the single-neuron frequency (**Fig. 3H’**) was detected. Together, these data suggest that the Munc18-1-mutant neurons exhibit a qualitative difference in network properties and in single-neuron Ca^2+^ dynamics. These results agree well with the previous finding that the heterozygous Munc18-1 deletion in human neurons causes a major decrease in synaptic strength^20^.

To validate the use of our analyses of the synchronous firing rate, we analysed the raw data by two currently available software as a comparison, FluoroSNNAP^12^ and PeakCaller^22^. Both of these ‘surrogate’ software quantified synchronization index as a measure of network activity. Analysis of our data using FluoroSNNAP (**Fig. 3E**) and PeakCaller (**Fig, 3F**) confirmed a decrease in the synchronization index in Munc18-1-mutant neurons compared to controls (**Fig. 3D**). Moreover, we also used the single-neuron module in FluoroSNNAP to quantify the Ca^2+^ signalling amplitude and frequency. Consistently, these analysis showed that Munc18-1-mutant neurons exhibit a reduction of the amplitude and an increase of the frequency compared to control neurons (**Fig. 3I & J**) without a difference in the coefficient of variation of the amplitude (**Fig. 3I’**) and frequency (**Fig. 3J’**). Together, these results validated the parameter quantifications we proposed with our approach.

### 3.5 Validation of the proposed Ca^2+^ imaging protocol using mouse neurons with a conditional deletion of all neurexins

In a second validation experiment, we examined cortical neurons cultured from newborn mice with a conditional deletion of all three neurexins, referred to as pan-neurexin deletion^13^. Neurexins are presynaptic adhesion molecules that act as key regulators of synapse properties^23^. Deletion of all neurexins has no effect on synapse numbers, but causes profound changes in the efficacy of synaptic transmission^13,24^. We infected the cultured neurons at DIV3 with lentiviruses expressing GCaMP6m and ΔCre (control) or Cre (deletion), and analysed them by Ca^2+^ imaging at DIV14-16 as described in section 2.2 (**Fig. 4A**).

**Fig. 4.**
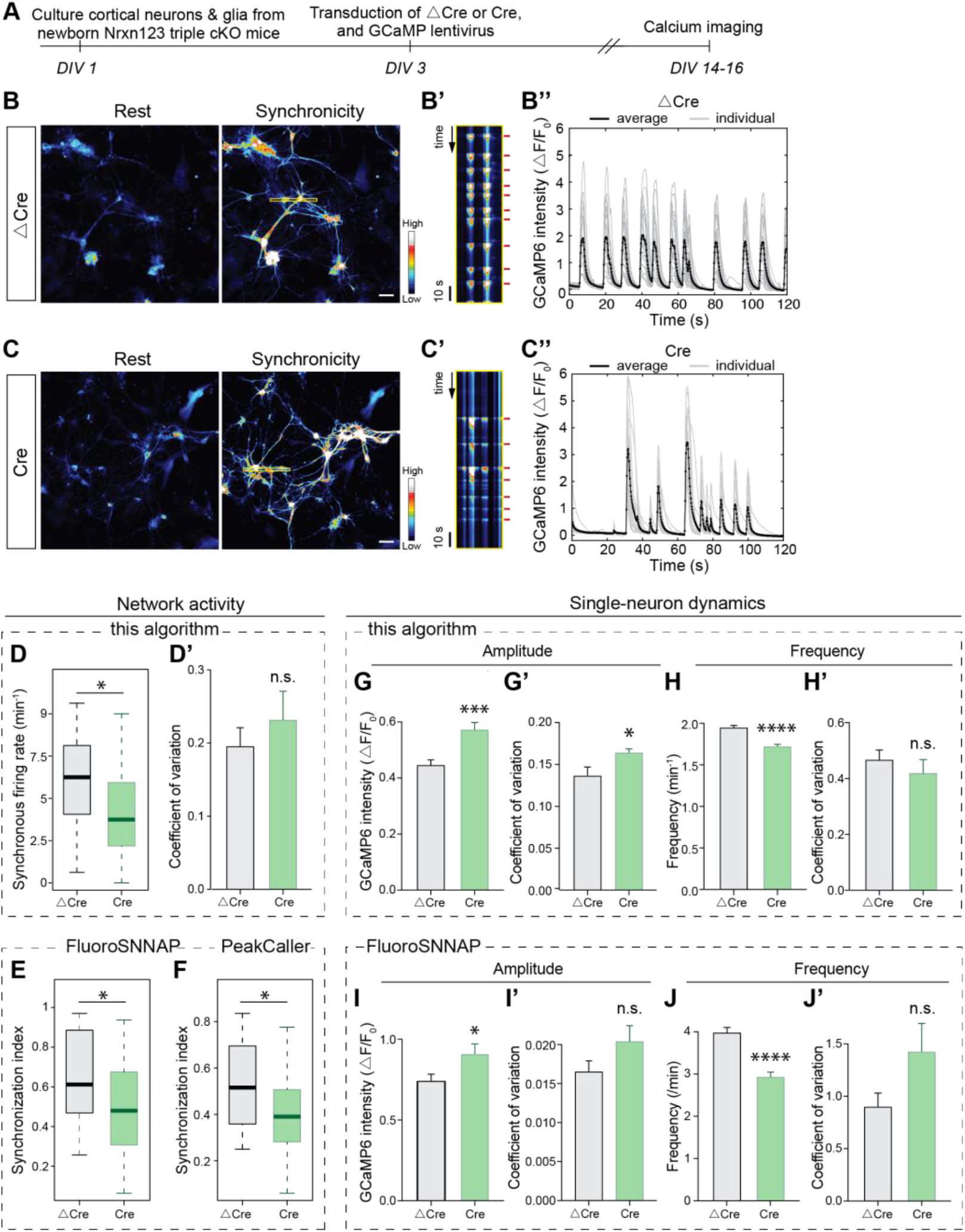
Analysis of network and single-neuron activities in cortical neurons cultured from triple neurexin-1/2/3 cKO mice. (**A**) Experimental strategy for culturing and imaging cortical neurons from triple neurexin-1/2/3 cKO mice. (**B-B’ & C-C’**) Ca^2+^ imaging analysis of network activity of control neurons (**B-B’**; triple neurexin-1/2/3 cKO neurons expressing ΔCre) and neurexin-deficient neurons (**C-C’**; triple neurexin-1/2/3 cKO neurons expressing Cre). Representative images of Ca^2+^ signals monitored in neurons in standard medium (Rest) and in medium containing 4 mM Ca^2+^ and 8 mM K^+^ (Synchronicity) are shown on the left (**B & C**); kymographs of the neurons included in the yellow boxes in the ‘Synchronicity’ images are depicted in the middle, with red lines marking Ca^2+^ spikes (right panels in **B & C**); and Ca^2+^ traces on the right (**B’ & C’**; thin lines depict individual neurons, and thick lines averages). (**D & D’**) Quantification of network activity using synchronicity firing rate (**D**) and the coefficient of variation of synchronous firing rate (**D’**) for control and triple neurexin-1/2/3 cKO neurons. (**E & F**) Quantification of synchronization index by FluoroSNNAP (**E**) and PeakCaller (**F**) for control and triple neurexin-1/2/3 cKO neurons. (**G & G’**) Quantification of single-neuron amplitude (**G**) and the coefficient of variation of amplitude (**G’**) for control and triple neurexin-1/2/3 cKO neurons using program proposed in this study. (**H & H’**) Quantification of single-neuron frequency (**H**) and the coefficient of variation of frequency (**G’**) for control and triple neurexin-1/2/3 cKO neurons using program proposed in this study. (**I & I’**) Quantification of single-neuron amplitude (**I**) and the coefficient of variation of amplitude (**I’**) for control and triple neurexin-1/2/3 cKO neurons using FluoroSNNAP. (**J & J’**) Quantification of single-neuron frequency (**H**) and the coefficient of variation of frequency (**G’**) for control and triple neurexin-1/2/3 cKO neurons using FluoroSNNAP. 890 control and 812 detected ROIs in neurexin-deficient neurons were used in **D-D’, F-H’**. 1476 control and 1282 detected ROIs in neurexin-deficient neurons were used in **E, I-J’**. 4 biological batches were used in **D-J’**. Data in **D-J’** represent means ± SEM. Student’s t-tests were used for all groups (*, p<0.05; ***, p<0.001; ****, p<0.0001; n.s., not significant). Scale bars, 25 μm. Objective, 20 X.

The pan-neurexin deletion significantly decreased neuronal network activity, as shown by a decreased synchronous firing rate (**Fig. 4B-D**). The reduced network activity in triple neurexin-1/2/3 cKO neuron cultures was further confirmed by the decrease of synchronization index measured by FluoroSNNAP (**Fig. 4E**) as well as by PeakCaller (**Fig. 4F**). At the single-neuron level, interestingly, the pan-neurexin deletion caused a significant, somewhat paradoxical increase of the amplitude of Ca^2+^ spikes (**Fig. 4G**), which was also confirmed by the single-neuron amplitude quantification using FluoroSNNAP (**Fig. 4I**). Opposite to the amplitude change, however, the frequency of Ca^2+^ spikes in pan-neurexin deletion neurons significantly decreased as quantified using our program (**Fig. 4H**) and using FluoroSNNAP (**Fig. 4J**). The variability of the network activity was examined by quantification of the coefficient of variation of synchronous firing rate, but no differences were observed between control and pan-neurexin deletion neurons (**Fig. 4D’**). The neuron-to-neuron variability was compared by quantification of the coefficient of variation of single-neuron amplitude and frequency using our program (**Fig. 4G’ & H’**) and using FluoroSNNAP (**Fig. 4I’ & J’**). Despite a slight increase of the coefficient of variation of the amplitude in the pan-neurexin deletion neurons measured by our program (**Fig. 4G’**), all other data detected little or no significant difference in pan-neurexin deletion neurons compared to controls (**Fig. 4H’, Fig. 4I’ & J’**).

## Discussion

In this study, we developed a Ca^2+^ imaging approach that enables economical and rapid functional analyses of the network activity and Ca^2+^ dynamics of cultured human and rodent neurons. The mechanisms underlying neuronal network activity have been studied extensively by patch-clamp electrophysiology, which provides an exquisite amount of precise information on individual neurons, but is only suitable for low-throughput applications. We demonstrated that the simple approach for the analysis of neuronal network activity and Ca^2+^ signalling that we describe yields reproducible, quantitative parameters. Among others, these parameters include the synchronicity of the network activity, and the amplitude and frequency of Ca^2+^ signals of individual neurons in larger populations. Since the neuronal network activity and the Ca^2+^ signals elicited by network activity depend on a large number of neuronal parameters, including the synaptic connectivity and the intrinsic electrical properties of neurons, the parameters that emerge from analyses of this network activity cannot be directly ascribed to a particular neuronal property, but are indicators of overall neuronal function that provide a useful measure for screening purposes.

In the approach proposed here, we induced synchronous network activity in cultured human and mouse neurons by raising the ambient Ca^2+^ and K^+^ concentrations, which enabled routine analyses of network activity without external stimulation. The increased Ca^2+^ and K^+^ concentrations stimulate network activity because they elevate the neurotransmitter release probability and the excitability of neurons. Although these conditions are not physiological, they trigger normal action potentials in cultured neurons, which in themselves are not a physiological preparation, but represent a reduced system amenable to mechanistic analyses. We validated our Ca^2+^-imaging approach with human and mouse neurons that carry defined conditional mutations in a synaptic gene, human heterozygous Munc18-1 and mouse homozygous pan-neurexin conditional KO neurons. The results we obtained with these mutant neurons confirmed that the simple Ca^2+^-imaging approach we describe is not only capable of robustly detecting mutant phenotypes, but also able to differentiate between phenotypes (**Figs. 3 and 4**). Moreover, we further validated our approach by analysing our Ca^2+^-imaging data obtained with the mutant neurons using two previously described software packages, FluoroSNNAP^12^ and PeakCaller^22^, demonstrating that our approach is likely more sensitive and versatile than these methods (**Figs. 3 and 4**; see also Table 1).

**Table 1.** Ca^2+^-imaging program feature comparisons and application scales.

| Study | Implementation | Features | Application scale |
| --- | --- | --- | --- |
| Patel et al., 2015<sup>12</sup> | Matlab (FluoroSNNAP) | Synchronization index<br>Network connectivity<br>Spike amplitude<br>Spike rise and decay time | Single or multiple neuron, *in vitro* mouse neuron. |
| Jee et al., 2015<sup>31</sup> | Matlab (NeuroCa)<br>[not open source] | Amplitude<br>Frequency<br>Background correction<br>Network dynamics | Single or multiple neuron, *in vitro* rat neuron. |
| Pnevmatikakis et al., 2016<sup>32</sup> | Matlab, Python | Cell and spike detection<br>Signal extraction, denoising and deconvolution | Single or multiple neuron, *in vivo* mouse neuron, zebrafish whole-brain, *in vitro* dendritic spine. |
| Artimovich et al., 2017<sup>22</sup> | Matlab (PeakCaller) | Synchronization index<br>Spike event number<br>Spike rise and fall time<br>Correlation coefficient | Single or multiple neuron, dendritic spines, *in vitro* mouse and human neuron. |
| Prada et al., 2018<sup>33</sup> | ImageJ, R (NA<sup>3</sup>) | x, y-field segmentation<br>Signal-to-noise ratio<br>Activity event counts | Single neuron, neurite activity, *in vitro* mouse motoneuron. |
| Mancini et al., 2018<sup>34</sup> | ImageJ, Matlab (SICT) | Frequency<br>Amplitude, integrated amplitude<br>Rise and fall time<br>FWHM | Single or multiple neuron, dendritic spines, *in vitro* mouse neuron. |
| Giovannucci et al., 2019<sup>27</sup> | Python, Matlab (CalmAn) | Large-scale, batch processing<br>Motion correction<br>Source extraction<br>Online processing | Single or multiple neuron, *in vivo* zebrafish whole brain light-sheet data. One-photon or two-photon, long-term recording. |
| This study | ImageJ, Matlab | Synchronization rate, amplitude<br>Coefficient of variation of synchronization rate<br>Spike amplitude, frequency<br>Coefficient of variation of spike amplitude, frequency | Single or multiple neuron, *in vitro* human and mouse neuron. |

In our approach, we adopted a semi-automated image analysis approach. For the manual part, users select interested neurons to be analysed (**Fig. 1B**), while the data extraction and computation of the synchronicity rate, single-neuron amplitude and frequencies, coefficients of variation are automatic. To test the robustness of the ROI selection as well as the parameter quantification methods in our protocol, we analysed our data with two independently available ‘surrogate’ software packages mentioned above, FluoroSNNAP^12^ and PeakCaller^22^. FluoroSNNAP provides automated ROI detection using time-lapse files as input and PeakCaller uses Ca^2+^ traces as input. Analysis of network activity as well as single-neuron dynamics using these software showed consistent differences between control and mutant neurons similar to those revealed by our approach (**Fig. 3D-J’, Fig. 4D-J’, Supplementary Fig. 1E-H, Supplementary Fig. 2E-H**). Additionally, because raw time-lapse movies were imported into FluoroSNNAP for analysis (**Supplementary Fig. 1A-B’, Supplementary Fig. 2A-B’**) while ROIs containing raw Ca^2+^ traces extracted using our programs were imported into PeakCaller for analysis (**Supplementary Fig. 1C-D, Supplementary Fig. 2C-D**), the consistent results of the synchronization index measured from the two software (**Fig. 3E & F; Fig. 4E & F**) suggested that the ROI detection method we proposed is robust. Overall, comparisons of the parameter values obtained (**Figs. 3 and 4**) suggest that our approach compares favourably with that of other software.

At present, Ca^2+^-imaging analysis packages are aiming to provide evaluations of diverse features at multiple levels and at different scales (see a partial list of available packages in **Table 1**)^12,26–29^. The increasing size of the imaging datasets demands that analyses should be fully automated, ultimately towards standardization and automation to leverage phenotypic assays especially in human neurons that can be used for drug screening purposes for neurodegenerative or neuropsychiatric disorders^1–3,5,30^. Our Ca^2+^ imaging approach is relatively simple compared to these more sophisticated packages. However, approach uses rather basic instrumentation and thus useful for non-industrial applications, and can be readily expanded to monitor additional parameters. For example, our protocol examines network activity as the frequency of synchronous peaks, yet in some cases the percentage of synchronously firing neurons might decrease without changes of the synchronous firing frequency^3^. To address this possibility, the algorithms we describe could be readily expanded to measure the percentage of neurons contributing to synchronous firing of the population. Furthermore, we quantify the Ca^2+^ signal from the soma of a neuron, which integrates the signals of hundreds of synaptic inputs. To measure Ca^2+^ signals directly at the level of synapses, GCaMPs could be targeted to pre- or to postsynaptic specializations by fusion to synaptophysin^25^ or to PSD95^11^, respectively. Thus the present study aims to enable a straightforward implementation of an important basic tool, measurements of Ca^2+^ signals in neuronal population, in order to facilitate widespread application of such measurements and to allow their expansion to other neuronal parameters as needed.

## Credit author statement

Z.S. and T.C.S. jointly conceived the experiments, Z.S. performed the experiments, and Z.S. and T.C.S. analysed the results and wrote the manuscript.

## Acknowledgements

The authors would like to thank Drs. Jinye Dai, Christopher Patzke, Steve Wilson, Kathlyn Gan, and Yang Yang for advice and reagents. This work was supported by a grant from the NIMH (U19 MH104172, to T.C.S.).

## SUPPLEMENTAL FIGURES and FIGURE LEGENDS

**Supplementary Figure 1:**
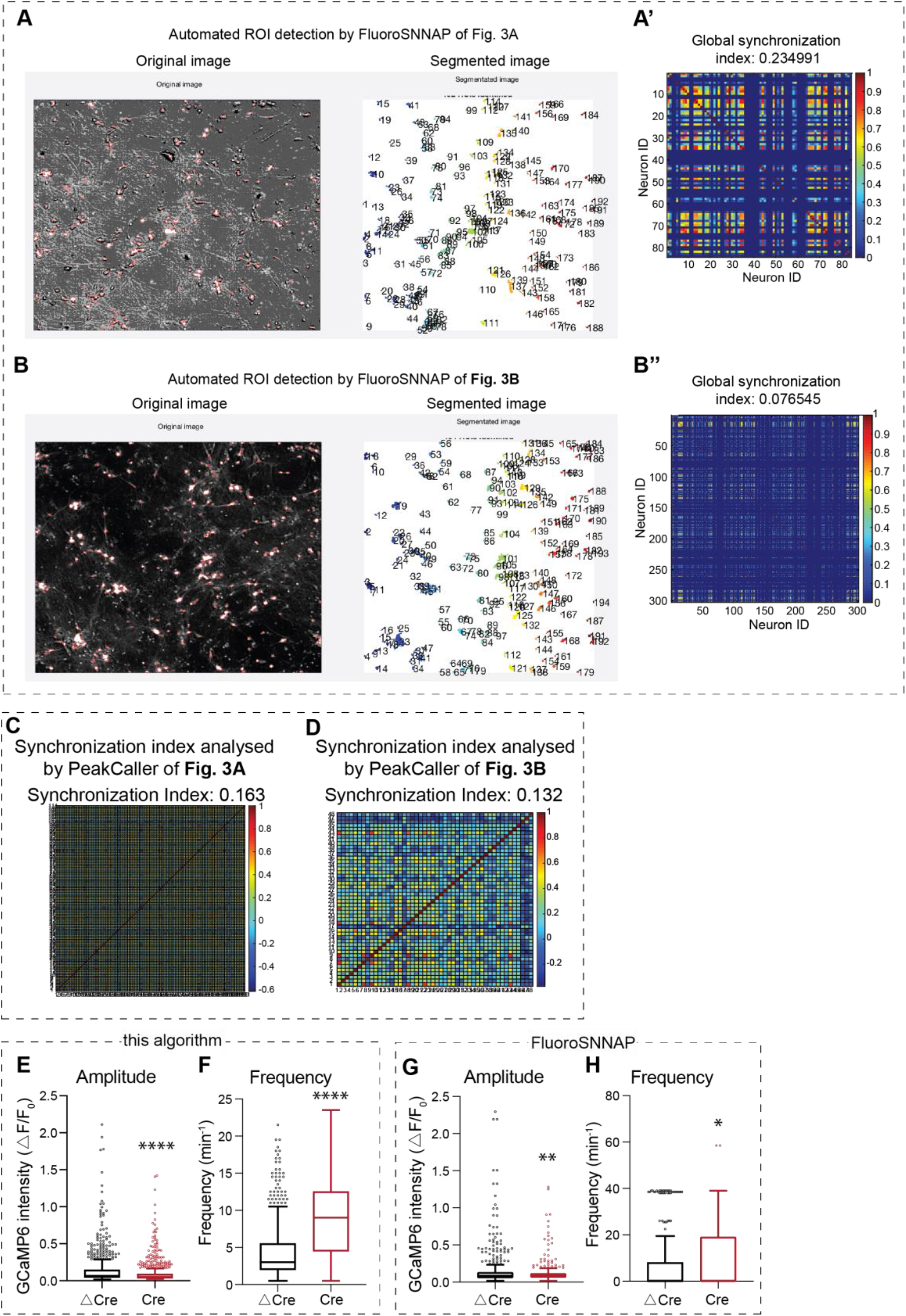
(**A & A’**) Example images showing the imported image and the segmentation panel using FluoroSNNAP of corresponding data in **Fig. 3B** (**A**), and the synchronization index quantification (**A’**). (**B & B’**) Images showing the imported image and the segmentation panel using FluoroSNNAP of corresponding data in **Fig. 3C** (**B**), and the synchronization index quantification (**B’**). (**C**) Example image showing the synchronization index quantified by PeakCaller of corresponding data in **Fig. 3B**. (**D**) Example image showing the synchronization index quantified by PeakCaller of corresponding data in **Fig. 3C**. (**E & F**) Box plot of quantification of single-neuron amplitude (**E**) and frequency (**F**) for control and heterozygous Munc18-1 mutant neurons using program in this study. The graphs correspond to the same data in **Fig. 3G & H**, respectively. (**G & H**) Box plot of quantification of single-neuron amplitude (**G**) and frequency (**H**) for control and heterozygous Munc18-1 mutant neurons using FluoroSNNAP. The graphs correspond to the same data in **Fig. 3I & J**, respectively. 689 and 981 neurons from control and Munc18-1-mutant neurons from 3 biological replicates were used in **E-F**. 1139 and 1274 detected ROIs from control and Munc18-1-mutant neurons from 3 biological replicates were used in **G-H**. Data in **E-H** represent mean ± SEM. Student’s *t*-tests were used for all groups (*, p<0.05; **, p<0.01; ****, p<0.0001). Objective, 10 X.

**Supplementary Figure 2:**
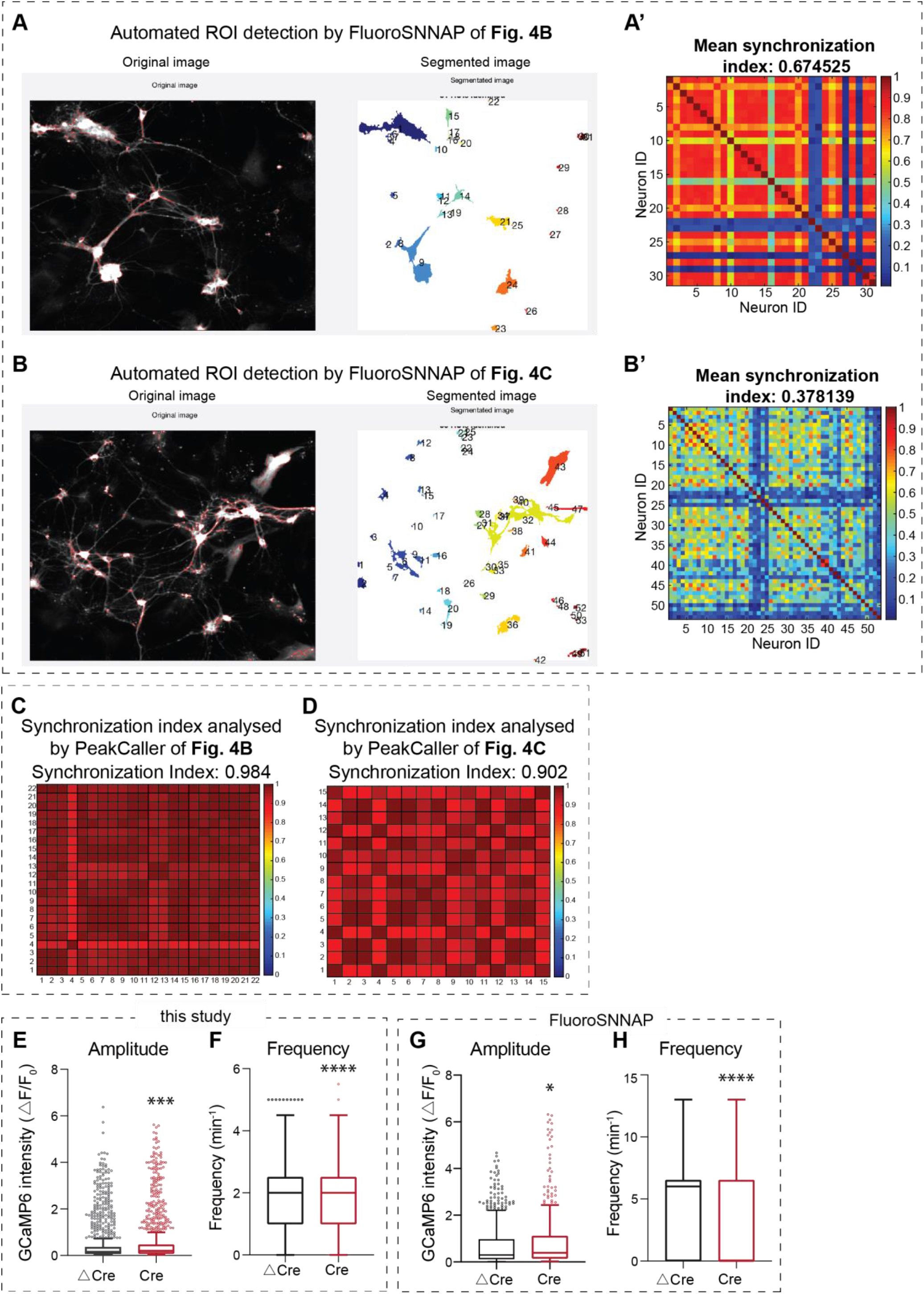
(**A & A’**) Example images showing the imported image and the segmentation panel using FluoroSNNAP of corresponding data in **Fig. 4B** (**A**), and the synchronization index quantification (**A’**). (**B & B’**) Example images showing the imported image and the segmentation panel using FluoroSNNAP of corresponding data in **Fig. 4C** (**B**), and the synchronization index quantification (**B’**). (**C**) Example image showing the synchronization index quantified by PeakCaller of corresponding data in **Fig. 4B-B’’**. (**D**) Example image showing the synchronization index quantified by PeakCaller of corresponding data in **Fig. 4C-C’’**. (**E & F**) Box plot of quantification of single-neuron amplitude (**E**) and frequency (**F**) for control and triple neurexin-1/2/3 cKO neurons using program in this study. The graphs correspond to the same data in **Fig. 4G & H**, respectively. (**G & H**) Box plot of quantification of single-neuron amplitude (**G**) and frequency (**H**) for control and triple neurexin-1/2/3 cKO neurons using FluoroSNNAP. The graphs correspond to the same data in **Fig. 4I & J**, respectively. 890 control and 812 detected ROIs in neurexin-deficient neurons from 4 biological batches were used in **G-H**. Data in **E-H** represent means ± SEM. Student’s t-tests were used for all groups (*, p<0.05; (*p<0.05; ***, p<0.001; ****, p<0.0001). Objective used, 20 X.

## References

1. Grienberger C, Konnerth A. Imaging calcium in neurons. Neuron 2012; 73(5): 862–85.

2. Verstraelen P, Pintelon I, Nuydens R, Cornelissen F, Meert T, Timmermans JP. Pharmacological characterization of cultivated neuronal networks: relevance to synaptogenesis and synaptic connectivity. Cell Mol Neurobiol 2014; 34(5): 757–76.

3. Verschuuren M, Verstraelen P, Garcia-Diaz Barriga G, et al. High-throughput microscopy exposes a pharmacological window in which dual leucine zipper kinase inhibition preserves neuronal network connectivity. Acta Neuropathol Commun 2019; 7(1): 93.

4. Brewer GJ, Boehler MD, Pearson RA, DeMaris AA, Ide AN, Wheeler BC. Neuron network activity scales exponentially with synapse density. J Neural Eng 2009; 6(1): 014001.

5. Verstraelen P, Van Dyck M, Verschuuren M, et al. Image-Based Profiling of Synaptic Connectivity in Primary Neuronal Cell Culture. Front Neurosci 2018; 12: 389.

6. Virdee JK, Saro G, Fouillet A, et al. A high-throughput model for investigating neuronal function and synaptic transmission in cultured neuronal networks. Sci Rep 2017; 7(1): 14498.

7. Wardill TJ, Chen TW, Schreiter ER, et al. A neuron-based screening platform for optimizing genetically-encoded calcium indicators. PLoS One 2013; 8(10): e77728.

8. Fan LZ, Nehme R, Adam Y, et al. All-optical synaptic electrophysiology probes mechanism of ketamine-induced disinhibition. Nat Methods 2018; 15(10): 823–31.

9. Williams LA, Joshi V, Murphy M, et al. Scalable Measurements of Intrinsic Excitability in Human iPS Cell-Derived Excitatory Neurons Using All-Optical Electrophysiology. Neurochem Res 2019; 44(3): 714–25.

10. Peled ES, Newman ZL, Isacoff EY. Evoked and spontaneous transmission favored by distinct sets of synapses. Curr Biol 2014; 24(5): 484–93.

11. Reese AL, Kavalali ET. Single synapse evaluation of the postsynaptic NMDA receptors targeted by evoked and spontaneous neurotransmission. Elife 2016; 5.

12. Patel TP, Man K, Firestein BL, Meaney DF. Automated quantification of neuronal networks and single-cell calcium dynamics using calcium imaging. J Neurosci Methods 2015; 243: 26–38.

13. Chen LY, Jiang M, Zhang B, Gokce O, Sudhof TC. Conditional Deletion of All Neurexins Defines Diversity of Essential Synaptic Organizer Functions for Neurexins. Neuron 2017; 94(3): 611–25 e4.

14. Smetters D, Majewska A, Yuste R. Detecting action potentials in neuronal populations with calcium imaging. Methods 1999; 18(2): 215–21.

15. Zhang Y, Pak C, Han Y, et al. Rapid single-step induction of functional neurons from human pluripotent stem cells. Neuron 2013; 78(5): 785–98.

16. Huang YA, Zhou B, Wernig M, Sudhof TC. ApoE2, ApoE3, and ApoE4 Differentially Stimulate APP Transcription and Abeta Secretion. Cell 2017; 168(3): 427–41 e21.

17. Huang YA, Zhou B, Nabet AM, Wernig M, Sudhof TC. Differential Signaling Mediated by ApoE2, ApoE3, and ApoE4 in Human Neurons Parallels Alzheimer’s Disease Risk. J Neurosci 2019; 39(37): 7408–27.

18. Tsetsenis T. Monitoring Synapses Via Trans-Synaptic GFP Complementation. Methods Mol Biol 2017; 1538: 45–52.

19. Piatkevich KD, Bensussen S, Tseng HA, et al. Population imaging of neural activity in awake behaving mice. Nature 2019; 574(7778): 413–7.

20. Patzke C, Han Y, Covy J, et al. Analysis of conditional heterozygous STXBP1 mutations in human neurons. J Clin Invest 2015; 125(9): 3560–71.

21. Sudhof TC. Neurotransmitter release: the last millisecond in the life of a synaptic vesicle. Neuron 2013; 80(3): 675–90.

22. Artimovich E, Jackson RK, Kilander MBC, Lin YC, Nestor MW. PeakCaller: an automated graphical interface for the quantification of intracellular calcium obtained by high-content screening. BMC Neurosci 2017; 18(1): 72.

23. Sudhof TC. Synaptic Neurexin Complexes: A Molecular Code for the Logic of Neural Circuits. Cell 2017; 171(4): 745–69.

24. Luo F, Sclip A, Jiang M, Sudhof TC. Neurexins cluster Ca(2+) channels within the presynaptic active zone. EMBO J 2020; 39(7): e103208.

25. Dreosti E, Odermatt B, Dorostkar MM, Lagnado L. A genetically encoded reporter of synaptic activity in vivo. Nat Methods 2009; 6(12): 883–9.

26. Kaifosh P, Zaremba JD, Danielson NB, Losonczy A. SIMA: Python software for analysis of dynamic fluorescence imaging data. Front Neuroinform 2014; 8: 80.

27. Giovannucci A, Friedrich J, Gunn P, et al. CaImAn an open source tool for scalable calcium imaging data analysis. Elife 2019; 8.

28. Oh J, Lee C, Kaang BK. Imaging and analysis of genetically encoded calcium indicators linking neural circuits and behaviors. Korean J Physiol Pharmacol 2019; 23(4): 237–49.

29. Zhou P, Resendez SL, Rodriguez-Romaguera J, et al. Efficient and accurate extraction of in vivo calcium signals from microendoscopic video data. Elife 2018; 7.

30. Mackay L, Mikolajewicz N, Komarova SV, Khadra A. Systematic Characterization of Dynamic Parameters of Intracellular Calcium Signals. Front Physiol 2016; 7: 525.

31. Jang MJ, Nam Y. NeuroCa: integrated framework for systematic analysis of spatiotemporal neuronal activity patterns from large-scale optical recording data. Neurophotonics 2015; 2(3): 035003.

32. Pnevmatikakis EA, Soudry D, Gao Y, et al. Simultaneous Denoising, Deconvolution, and Demixing of Calcium Imaging Data. Neuron 2016; 89(2): 285–99.

33. Prada J, Sasi M, Martin C, Jablonka S, Dandekar T, Blum R. An open source tool for automatic spatiotemporal assessment of calcium transients and local ‘signal-close-to-noise’ activity in calcium imaging data. PLoS Comput Biol 2018; 14(3): e1006054.

34. Mancini R, van der Bijl T, Bourgeois-Jaarsma Q, Lasabuda R, Groffen AJ. SICT: automated detection and supervised inspection of fast Ca(2+) transients. Sci Rep 2018; 8(1): 15523.

