## Appendix A for "A simple Ca^2+^-imaging approach to neural network analysis in cultured neurons"

**Analysis of calcium dynamics**

i. **CONTENTS**
ii. **i. Introduction**
iii. **ii. Installation and file preparation**
iv. **iii. Results and Interpretation**
v. **iv. Troubleshooting**

**I Introduction**

**——————————————————————————————————————**

This version of the Matlab-based analysis program package contains five functions / scripts:

1. GCaMP_multiROI.m defines ROIs, and calculates the intensity of selected ROIs.
2. Raw_Intensity.m plots the raw intensity of all calcium traces.
3. Synchronous_rate_amplitude.m calculate the synchronous firing rate & amplitude from the average intensity of all ROIs in one FOV.
4. SingleNeuron_Amplitude_ddF0_Frequency.m firstly detect spikes, normalises the raw intensity to deltaF/F0, calculates the amplitude profile at individual cell level and quantify all cells. Secondly, the script also quantifies the frequency of spikes (number of detected spikes / min). At last, the script also compute the coefficient of variation of the amplitude and frequency.

**II Installation and File preparation**

**——————————————————————————————————————**

II-a Installation

Please follow the instruction to download and install Matlab.

II-b File preparation

The live-imaging image stack should be converted to TIFF/TIF format for Matlab analysis. You can do this either by exporting the file through the microscope analysis software, for example if using Nikon microscope it is the NIS; or you can use ImageJ (or FIJI) to export the nd2 format to TIFF/TIF format.

**III Results and Interpretation**

**——————————————————————————————————————**

**III-a GCAMP_multiROI.m**

Note the variables in the function are:

- timerinterval = the frame rate in second.
- name1 = the COMPLETE file name of the tiff image stack you are going to analyse.
- legend1 = the figure legend you want to put.
- diameter = the diameter of ROI in micrometer - it is roughly the cell diameter.
- varargin = leave this blank as it will automatically save all the parameters in a matrix.

To run the "GCAMP_multiROI.m" function, see an example here:

to analyse a file named 'neuron.tif' which has time interval of 20 msec and the neuron soma diameter is 6um, the function should be:

GCAMP_1_multiROIplot(0.02,'neuron.tif’,’GCaMP intensity',6).

Upon running the function, it takes 10-30 sec to prompt an image which is the maximum intensity projection of the image stack, then select the ROI/cells on this image you want to analyse, then press Enter key to finish all the selections.

The function will calculate intensity profile of each cells and its coordinate all saved in the file named ROI_intensity1.mat. Each column is one cell, and each the row is the intensity at each time point.

You can export the data into Excel, csv, txt etc for further analyses.

**III-b Raw_Intensity.m**

This is a simple script to plot all the original calcium traces as well as the averaged intensity.

To use it:

Load your data first - load the matrix named 'ROI_intensity1' generated from above step or other matrix, txt file.

Then click 'Run' button in the script window, or copy & paste the entire script to the Command region and press Enter key.

**III-c Synchronicity_rate_amplitude.m**

This is a simple script to calculate the synchronous firing rate & amplitude from the averaged intensity of all ROIs.

To use it:

Load your data first - load the matrix named 'ROI_intensity1' generated from above step or other matrix, txt file.

Then click 'Run' button in the script window, or copy & paste the entire script to the Command region and press Enter key.

The synchronous firing rate is defined as = number of synchronous spikes detected in the average intensity of all ROI / min, which is saved in Sync_rate.

The synchronous firing amplitude is defined as = deltaF to F value of synchronous spikes detected in the average intensity of all ROI / min, which is saved in Sync_amplitude_dFF.

Both value will be displayed in the command window after run the script. And the outputs can be found in the matrix:

Sync_rate: the synchronous firing rate in this FOV

Sync_dFF = the synchronous firing amplitude in this FOV

cov_Sync_dFF = the coefficient of variation of synchronicity (measured by the standard deviation of synchronous firing amplitude / mean of synchronous firing amplitude)

**III-d SingleNeuron_Amplitude_ddF0_Frequency.m**

This is the script to plot the normalized calcium intensity as well as to extract the amplitude data.

To use it:

Load your data (matrix, txt file) first, then:

simply click 'Run' button in the Matlab working environment, or copy & paste the entire script to the Command region and press Enter key.

This will return the outputs:

- Two plots: the plot of detected spikes and the plot of normalized calcium traces.
- In the dFF0 array, it contains the averaged amplitude from the detected spikes of individual cell;
- In the deltaFtoF0 matrix contains the value of normalized intensity dF/F0 as a function of time of all cells.
- In the neuron_freq, it contains the the frequency of spikes (detected spikes / min) for each neuron.
- In the std_dff, it contains the standard deviation quantification of amplitude (dff) for each neuron.
- In the std_freq, it contains the standard deviation quantification of frequency for each neuron.
- In the cov_dff, it contains the coefficient of variation of amplitude (dff).
- In the cov_freq, it contains the coefficient of variation of frequency.

*****NOTE** Adjust the time interval, recording duration accordingly

*****NOTE** Adjust the X-/Y-axis limit according to each plot or the specifications of the curve as needed.

*****NOTE** Adjust the threshold if needed, normally it should be mean + 1-2.5* std.

**IV Troubleshooting**

**——————————————————————————————————————**

This analysis package provides essential command and is complete for the above described analysis purpose, but can be optimized and modified.

If the program does not execute, check the following first:

whether the Matlab function files are placed in the same saving path where you save your data. if not, you should click the 'Add to path' -> 'Selected folder and subfolders'.

Comments and reports of bugs can be sent to:

——————————————————————————————————————
