## Appendix B1 for "A simple Ca^2+^-imaging approach to neural network analysis in cultured neurons"

```
function [ROI_intensity1, c, r]=GCAMP_multiROIonePLOT(timeinterval,name1,legend1,diameter,varargin)

%% plot 2 wavelength together with specific time axis
%name1 in FULL file name, second in red,pick rigion from name1
%timeinterval=3;
%diameter=10; % 2*diameter+1 is the square used for zoom in ROI
%plot is normalized to 0-1, and .mat file save the original intensity
%numbers.
%plot and .mat file will be saved in a new folder with the same as the movie data.

line_width=0.5;
marker_size=0.5;
tic;
saveinfo=strrep(name1,'.tif','_multiROIonePLOT/');
mkdir(saveinfo);
[~,savename1,~]=fileparts(name1);

%% load image
InfoImage=imfinfo(name1);
WImage=InfoImage(1).Width;
HImage=InfoImage(1).Height;
NumberImages=length(InfoImage);
ImageStack1=zeros(HImage,WImage,NumberImages,'uint16');

[t]=(0:NumberImages);
v=zeros(NumberImages,1);
for i=1:NumberImages
    v(i)=t(i)*timeinterval;
end
%current_t = clock;

for n=1:NumberImages
    ImageStack1(:,:,n)=imread(name1,'Index',n);
end

toc;
%% calculate median of 10-40 of the first 50 frames for preview

median_image=uint16(median(double(ImageStack1(:,:,10:40)),3));


%%  pick any region of interest

if ~isempty(varargin);
    c=varargin{1};
    r=varargin{2};
else
    scale = stretchlim(median_image);
    scrsz = get(0,'ScreenSize');
    figure('Position',[scrsz(3)*0.8 scrsz(4)*0.35 WImage HImage]);
    set(gca,'Position',[0 0 1 1]);
    imshow(imadjust(median_image,scale));
    [c, r, ~]= impixel;% select pixel
end

 colors=lines(length(c));

%% quantify multiple region intensity
ROI_intensity1=zeros(NumberImages,length(r));

cd(saveinfo);

handle=figure('PaperPosition',[0.25 2.5 2.4 0.2],'PaperUnits', 'inches');
for k=1:length(r)
    ROI_intensity1(:,k)=mean(mean(ImageStack1(max(1,r(k)-diameter):min(r(k)+diameter,HImage),max(1,c(k)-diameter):min(c(k)+diameter,WImage), :)));
    %transient=(ROI_intensity1(:,k)-min(ROI_intensity1(:,k)))/(max(ROI_intensity1(:,k))-min(ROI_intensity1(:,k)));
    plot(v(1:NumberImages),ROI_intensity1(:,k),'-o','Color',colors(k,:),'LineWidth',line_width,...
        'MarkerSize',marker_size);
    % print('-dtiff','-r150', [name1 '_median_d' num2str(diameter) '_c' num2str(c(k)) '_r' num2str(r(k)) '.tiff']);
    hold on;
end
    %xlimit=timeinterval*NumberImages;
    xlimit=50;
    yl = ylim;
    set(gca,'XLim',[0 xlimit],'YLim',yl,'LineWidth',1,'fontsize',8,'Fontname','Arial');
    ax = gca;
    ax.XGrid = 'on';
    ax.YGrid = 'off';
    ax.GridLineStyle = ':';
    %set(gca,'YLim',[-0.2 1.5],'LineWidth',1);
    box off;
    xlabel('time/s','Fontname','Arial');
    ylabel('intensity','Fontname','Arial');
    % title('IgE internalization after stimulation','fontsize',18,'fontweight','b','color','blue')legend(legend1,legend2,'Location','Northoutside','fontSize',8,'Plotboxaspectratio',[0.5 0.5 0.5]);
   % hl=legend('cell1','cell2','cell3','Location','NorthEast','Orientation','horizontal');
    %hl=legend('cell1','cell2','cell3','Location','NorthEast');
   % rect = [0.7, 0.7, .25, .3];
    %set(hl, 'Position', rect)
    %legend('boxoff');
    %h=legend(h,'KD','WT','Location','NorthWest');
    set(findall(handle,'type','text'),'fontSize',8,'LineWidth',2);
    % print('-dtiff','-r150', [saveinfo '2cplot_d' num2str(diameter) '_c' num2str(c(k)) '_r' num2str(r(k)) '.tiff']);
%    print('-depsc','-r150', [saveinfo '2cplot_d' num2str(diameter) '_c' num2str(c(k)) '_r' num2str(r(k)) '.eps']);
    hold off;

    %plotyy(v(1:201,2),ROI_intensity(:,k),v(1:201,2),ROI_intensity(:,k));

    close all;


%% save mat file   %temperally % .mat saving
save([savename1 '_median_d' num2str(diameter) '_c' num2str(c(1)) '_r' num2str(r(1)) '_plot.mat'],'ROI_intensity1','c','r','diameter','name1','timeinterval');
%% display the first frame overlaid with ROI
FirstImage=ImageStack1(:,:,1);
crop_width=diameter;
crop_height=diameter;
colors=autumn(length(c));

scrsz = get(0,'ScreenSize');
resolution=get(0,'ScreenPixelsPerInch'); % mine is 72
figure('Position',[scrsz(3)*0 scrsz(4)*0.1 WImage HImage],...
    'PaperPosition',[0.25 2.5 WImage/resolution HImage/resolution],...
    'PaperUnits', 'inches');% have to use screen resolution with combined lineart to make sure it keeps the same ratio
set(gca,'Position',[0 0 1 1]);
scale = stretchlim(FirstImage,[0.01 0.999]);
imshow(imadjust(FirstImage,scale));
axis off,
set(gca,'XTick',nan,'YTick',nan);
hold on;
for k=1:length(r)

    cidx = mod(k,length(c))+1;
    %rectangle('Position',[c(k)-crop_width,r(k)-crop_height,2*crop_width+1,2*crop_height+1],'EdgeColor',colors(cidx,:),'LineWidth',4); % rectangle ROI
    rectangle('Position',[c(k)-crop_width,r(k)-crop_height,2*crop_width+1,2*crop_height+1],'Curvature',[1 1],'EdgeColor',colors(cidx,:),'LineWidth',3); % round ROI
    hold on;
   % h = text(c(k),r(k), [num2str(c(k)) ',' num2str(r(k))]);
    %set(h,'Color',colors(cidx,:),'FontSize',15,'Fontname','Arial');
end
hold off;
print('-dtiff',[savename1 '_d' num2str(diameter) '_c' num2str(c(1)) '_r' num2str(r(1)) '_1cplot.tiff']);close all;


%% save median file
save([savename1 '_d' num2str(diameter) '_1cplot.mat'],'ROI_intensity1','c','r','diameter','name1');

%% display image overlaid with ROI square

FirstImage=zeros(HImage,WImage,1,'uint16');
FirstImage(:,:,1)=imread(name1,'Index',1);
crop_width=diameter;
crop_height=diameter;
colors=autumn(length(c));

% figure('PaperPosition',[0.25 2.5 6 4],'PaperUnits', 'inches');
scrsz = get(0,'ScreenSize');
resolution=get(0,'ScreenPixelsPerInch');
figure('Position',[scrsz(3)*0 scrsz(4)*0.1 WImage HImage],...
    'PaperPosition',[0.25 2.5 WImage/resolution HImage/resolution],...
    'PaperUnits', 'inches');% have to use screen resolution with combined lineart to make sure it keeps the same ratio
set(gca,'Position',[0 0 1 1]);
scale = stretchlim(FirstImage,[0.01 0.999]);
imshow(imadjust(FirstImage,scale));
axis off,
set(gca,'XTick',nan,'YTick',nan);
hold on;
for k=1:length(r)
    cidx = mod(k,length(c))+1;
    % rectangle('Position',[c(k)-crop_width,r(k)-crop_height,2*crop_width+1,2*crop_height+1],'EdgeColor',colors(cidx,:),'LineWidth',4);% rectangle ROI
    rectangle('Position',[c(k)-crop_width,r(k)-crop_height,2*crop_width+1,2*crop_height+1],'Curvature',[1 1],'EdgeColor',colors(cidx,:),'LineWidth',3); % round ROI
    hold on;
    %h = text(c(k),r(k), [num2str(c(k)) ',' num2str(r(k))]);
    %set(h,'Color',colors(cidx,:),'FontSize',15,'Fontname','Arial','FontWeight','bold');
end
hold off;
%cd(dir);
print('-dtiff',[ savename1 '_d' num2str(diameter) '_1cplot.tiff']);
close all;

%%
toc;

close all;
```
