## Appendix B2 for "A simple Ca^2+^-imaging approach to neural network analysis in cultured neurons"

%% Plot raw intensity

handle=figure('PaperPosition',[0.25 2.5 2.5 1.2],'PaperUnits', 'inches');

lclr=[0.6,0.8,1];

clr=[0.2,0,1];

b=1:length(ROI_intensity1(:,1));

timeinterval=0.0545; %define time resolution in sec

t=b*timeinterval;

figure;plot(t,ROI_intensity1,'LineWidth',0.5,'LineStyle',':');

hold on;

avg=mean(ROI_intensity1');

plot(t,avg,'Color','k','LineWidth',2);

xlim([0 150])

ylabel('GCaMP6 intensity / a.u.','Fontname','Arial','FontSize', 14);

xlabel('Time / sec', 'Fontname','Arial','FontSize', 14);
