## Appendix B3 for "A simple Ca^2+^-imaging approach to neural network analysis in cultured neurons"

```
%% define the average intensity into matrix
data=mean(ROI_intensity1');
timeinterval=0.054446; % Define recording time resolution in second, example here 0.0545 sec
rec_duration = 2.5; % Define recording duration in minute, example here 2.5min
[num_record,data_len]=size(data);
amplitudes = cell(num_record,1);

%% calculate the amplitude of major spikes above threshold
for j=1:num_record
    spike_data = data(j,:);
    sigma = std(data(j,:));
    threshold = mean(data(j,:)) + 1.5 * sigma; % Adujust the threshold based on the variation of the data, should be within 2-4 * sigma
    spike_index = spike_detection(spike_data, threshold);
    amplitude = data(j, spike_index);
    timestamp{j} = sparse(j,spike_index,amplitude);
    amplitudes{j} = amplitude;
end

%% calculate ampltiude, dff amplitude of major synchronous spikes
mid=round(data_len/2);
for m = 1:num_record
    min1 = min(data(m,1:100));
    min2 = min(data(m,(mid-50):(mid+50)));
    min3 = min(data(m,(data_len-100):(data_len)));
    mins = [min1 min2 min3];
    F0(m,1) = mean(mins);
    amplitudes_dff{m} = (amplitudes{m} - F0(m,1))./F0(m,1);
    mean_Sync_amplitude_dFF(m,1) = mean((amplitudes{m} - F0(m,1))./F0(m,1));
end
std_Sync_dff = std(amplitudes_dff{1,:});
cov_Sync_dff = std(amplitudes_dff{1,:})./ mean_Sync_amplitude_dFF;
Sync_dFF = amplitudes_dff{1,1}';

%% calculate rate of synchronous peaks
[zz_reco, num_SynPeaks] = size(amplitudes{1,1});
Sync_rate= num_SynPeaks / rec_duration;

%% plot deconvoluted spikes into cell structure & display values
delta = 1;
hold on;
for i = 1:num_record
    plot((num_record - i)*delta + timestamp{i}(i,:), '-', 'linewidth', 1);
end
set(gca, 'YTick', [], 'ycolor', 'k');
ylabel('Synchronous spikes detedted','Fontname','Arial', 'FontSize', 16);
xlabel('Time / frame','Fontname','Arial', 'FontSize', 16);

%Display synchronous rate & amplitude
display(['Synchronous rate =',num2str(Sync_rate(1,1)),'; Synchronous amplitude (dFF0)=',num2str(mean_Sync_amplitude_dFF(1,1))]);

%% plot raw intensity to check the synchronous peaks in raw plot (optional)
handle=figure('PaperPosition',[0.25 2.5 2.5 1.2],'PaperUnits', 'inches');
lclr=[0.6,0.8,1];
clr=[0.2,0,1];
b=1:length(ROI_intensity1(:,1));
timeinterval=0.054585; %define time resolution in sec
t=b*timeinterval;
figure;plot(t,ROI_intensity1,'LineWidth',0.5,'LineStyle',':');
hold on;
plot(t,data,'Color','k','LineWidth',2);
xlim([0 150]) % Define recording duration in sec, example here 150sec
ylabel('GCaMP6 intensity / a.u.','Fontname','Arial','FontSize', 14);
xlabel('Time / sec', 'Fontname','Arial','FontSize', 14);
```
