## Appendix B4 for "A simple Ca^2+^-imaging approach to neural network analysis in cultured neurons"

```
%% define data matrix
data=ROI_intensity1';
[num_record,data_len]=size(data);
amplitudes = cell(num_record,1);

%% Calculate the amplitude of all peaks above threshold
for j=1:num_record
    spike_data = data(j,:);
    sigma = std(data(j,:));
    threshold = mean(data(j,:)) + 1.5 * sigma;  %Adujust the threshold based on the variation of the data, should be mean + 1-2 *std
   % threshold = 2 * sigma; % Second way of setting threshold, in this way the threshold is 2-4 * sigma
    spike_index = spike_detection(spike_data, threshold);
    amplitude = data(j, spike_index);
    timestamp{j} = sparse(j,spike_index,amplitude);
    amplitudes{j} = amplitude;
end
%% To plot the un-normalized amplitude into the cell structure
delta = 1;
hold on;
for i = 1:num_record
    plot((num_record - i)*delta + timestamp{i}(i,:), '-', 'linewidth', 1);
end
%ylim([0,200000]) %Adjust the Y axis limit based on each data
set(gca, 'YTick', [], 'ycolor', 'k');
ylabel('Spike detedted in trace', 'FontSize', 14);
xlabel('Time / frame', 'FontSize', 14);

%% calculate baseline F0, deltaF/F0, std of dFF0 for each neuron
mid=round(data_len/2);
for m = 1:num_record
    min1 = min(data(m,1:100));
    min2 = min(data(m,(mid-50):(mid+50)));
    min3 = min(data(m,(data_len-100):(data_len)));
    mins = [min1 min2 min3];
    F0(m,1) = mean(mins);
    amplitudes_dff{m} = (amplitudes{m} - F0(m,1))./F0(m,1); %normalize to dff for individual neuron
    deltaFtoF0(m,1) = mean(amplitudes_dff{m});
end
ave_dff = mean(deltaFtoF0);
std_dff = std(deltaFtoF0);
cov_dff = std(deltaFtoF0)./mean(deltaFtoF0); % calculate the coeffiecient of variation of amplitude
%% calculate number, freq of spikes, std of freq of individual neuron
for k = 1:num_record
    [zz_zero, num_spikes{k}] = size(amplitudes{k});
    neuron_freq(k,1) = num_spikes{k} / 5; % the frequency is number of spikes/ min, define the recording duration, here is 2min
end
    ave_freq = mean(neuron_freq);
    std_freq = std(neuron_freq); %standard deviation of frequency
    cov_freq = std(neuron_freq)./ mean(neuron_freq); % coefficient of variability of single-neuron frequency
%% plot the deltaF/F0 in black & gray
handle=figure('PaperPosition',[0.25 2.5 2.5 1.2],'PaperUnits', 'inches');
reflclr=[0.8,0.8,0.8];
refclr='k';
F0plot=F0';
for i=1:num_record
    dFF0(:,i) = (ROI_intensity1(:,i)-F0plot(:,i))./F0plot(:,i);
end
b=1:data_len;
timeinterval=0.054585; %Adjust timeinterval, the unit is in sec
t=b*timeinterval;
plot(t,dFF0,'Color',reflclr,'LineWidth',0.5,'LineStyle','-');hold on;
avg=mean(dFF0');
plot(t,avg,'-o','Color',refclr,'LineWidth',2,'MarkerSize',2);
ylim([-1 2.2])
xlim([0 150]) %Adjust the recording duration, here it is 150 sec
ylabel('deltaF/F0 of GCaMP6 intensity', 'Fontname','Arial', 'FontSize', 14);
xlabel('Time / sec', 'Fontname','Arial','FontSize', 14);

% close all

%% plot the deltaF/F0 in random color
% handle=figure('PaperPosition',[0.25 2.5 2.5 1.2],'PaperUnits', 'inches');
% % reflclr=[0.8,0.8,0.8];
% % refclr='k';
% F0plot=F0';
% for i=1:num_record
%     dFF0(:,i) = (ROI_intensity1(:,i)-F0plot(:,i))./F0plot(:,i);
% end
% b=1:data_len;
% timeinterval=0.054585; %Adjust timeinterval, the unit is in sec
% t=b*timeinterval;
% plot(t,dFF0,'LineWidth',0.4,'LineStyle','-');hold on;
% avg=mean(dFF0');
% plot(t,avg,'-o','Color','k','LineWidth',2,'MarkerSize',2);
% ylim([-1 2.2])
% xlim([0 150]) %Adjust the recording duration, here it is 150sec
% ylabel('deltaF/F0 of GCaMP6 intensity', 'Fontname','Arial', 'FontSize', 14);
% xlabel('Time / sec', 'Fontname','Arial','FontSize', 14);
% %close all
```
